# Robust and replicable measurement for prepulse inhibition of the acoustic startle response

**DOI:** 10.1101/601500

**Authors:** Eric A. Miller, David B. Kastner, Michael N. Grzybowski, Melinda R. Dwinell, Aron M. Geurts, Loren M. Frank

## Abstract

Measuring animal behavior in the context of experimental manipulation is critical for modeling and understanding neuro-psychiatric disease. Prepulse inhibition of the acoustic startle response (PPI) is a behavioral paradigm used extensively for this purpose, but the results of PPI studies are often inconsistent. As a result, the utility of this metric remains uncertain. Here we deconstruct the phenomenon of PPI. We first confirm several limitations of the traditional PPI metric, including that the underlying startle response has a non-Gaussian distribution and that the traditional PPI metric changes with different stimulus condition. We then develop a novel model that reveals PPI to be a combination of the previously appreciated scaling of the startle response, as well as a scaling of sound perception. Using our model, we find no evidence for differences in PPI in a rat model of Fragile-X Syndrome (FXS) compared to wild-type controls. These results in the rat provide a reliable methodology that could be used to clarify inconsistent PPI results in mice and humans. In addition, we find robust differences between wild-type male and female rats. Our model allows us to understand the nature of these differences, and we find that both the startle-scaling and sound-scaling components of PPI are a function of the baseline startle response. Males and females differ specifically in the startle-scaling, but not the sound-scaling, component of PPI. These findings establish a robust experimental and analytical approach that has the potential to provide a consistent biomarker of brain function.

## Introduction

Prepulse inhibition of the acoustic startle response (PPI) is a reduction in the magnitude of the acoustic startle response when a weak, non-startling sound—the prepulse—precedes an intense, potentially-startling, sound^1–3^. Changes in PPI have been linked to various neuropsychiatric disorders, such as schizophrenia^4–9^, obsessive compulsive disorder^10–12^, Tourette’s syndrome^13,14^, autism spectrum disorder^15–17^, and posttraumatic stress disorder^18,19^. As such, PPI has been promoted as a potential biomarker of brain function in the context of disease^20,21^. Furthermore, since PPI can be studied in both humans and laboratory animals, it offers a translational methodology for generating mechanistic insights into those diseases^22–24^.

However, published PPI results are often inconsistent with one another^25^, potentially undermining the utility of the measure. The source of these inconsistencies has been associated with differences between experimental conditions^26^, analytical methods^24^ or factors such as strain^27,28^, age^29–31^, sex^32–35^, reproductive cycle^36,37^, species^38–40^, acute disease state^41^, habituation^42^, socialization^43,44^, and the baseline startle response^45^. Consequently, there is a pressing need for an approach that could consistently identify real differences among groups. We therefore sought to deconstruct the phenomenon of PPI to develop a more accurate methodology for capturing the way in which a prepulse stimulus modifies the acoustic startle response.

The traditional methodology for PPI makes four assumptions: 1) the startle response can be accurately measured with a small number of trials per animal; 2) the startle response has an approximately Gaussian distribution, allowing the use of the mean startle response as the basis of the PPI metric; 3) PPI is stable across startle sound levels, enabling the measurement of PPI at a single startle level instead of necessitating a full measurement of the startle function; and 4) PPI is independent of the baseline startle response, allowing for a direct combination of PPI results between animals.

Using data from 72 rats across more than 100 stimulus conditions, for a total of over 300,000 trials, we replicated previous work demonstrating that the aforementioned assumptions do not hold. Specifically, our findings confirm: 1) the startle response is highly variable^46^; 2) the startle response has a non-Gaussian distribution that is better represented by a log-normal distribution^47^; 3) the traditional metric used for PPI systematically decreases as a function of sound level^48^; and 4) PPI is also a function of the baseline startle response^45^.

These problems were identified in individual studies, but a systematic approach to account for all of them is lacking. Therefore, we present a novel analytical model of PPI characterized by a functional scaling of both the startle response and the startle sound. We fit a single model to these two components and found that it better fits the data than the implicit model underlying the traditional PPI metric.

Using our model, and data from multiple cohorts of animals, we conclude that *Fmr1* knockout (KO) rats—rats missing the gene silenced in Fragile-X Syndrome (FXS)—do not differ from wild-type (WT) rats in PPI. In contrast, we found that WT female rats differ from WT male rats in the startle-scaling, but not the sound-scaling, component of PPI. These experimental findings demonstrate the utility of our approach to generate robust and replicable PPI results. This approach, grounded in a formal mathematical model, has the potential to yield consistent findings about the relationship between PPI and genetic or experimental manipulations. As such, this approach could be used to clarify the inconsistent PPI results in the context of brain diseases, such as those reported in mouse models of Fragile-X Syndrome^39,49–56^.

## Materials and methods

### Animals

All experiments were conducted in accordance with Medical College of Wisconsin and University of California San Francisco Institutional Animal Care and Use Committee and US National Institutes of Health guidelines. Rat datasets were collected from Long Evans rats that were fed standard rat chow (LabDiet 5001).

The *Fmr1* KO rats were males with a CRISPR/SpCas9 knockout of *Fmr1* on a Long Evans background generated at the Medical College of Wisconsin. Briefly, a CRISPR targeting the *Fmr1* exon 8 sequence 5’-GGTCTAGCTATTGGTACTCA**TGG**-3’ (PAM in bold) was injected into Crl:LE embryos (Charles River Laboratories). Two mutant strains were generated (LE-*Fmr1*^*em2Mcwi*^ and LE*-Fmr1*^*em4Mcwi*^ (RGDIDs: 11553873, 11553875) with mutations in *Fmr1*. LE-*Fmr1*^*em2Mcwi*^ harbors a net 2-bp insertion, while LE*-Fmr1*^*em4Mcwi*^ harbors a 2-bp deletion mutation at the SpCas9 cleavage site (Fig. S1a). Both mutations are predicted to cause frame-shifts and complete loss of FMR1 expression was confirmed by Western blot (data no shown). Like *Fmr1* knockout mice^57^, knockout rats tended to be slightly larger and had significantly increased testicular weights at 30 days of age (Fig. S1b). Breeding colonies for both strains are maintained at the Medical College of Wisconsin in continuous backcross of heterozygous females to vendor Crl:LE males at each generation to avoid inbreeding and genetic drift.

There was a total of 72 rats in five different cohorts. The first *Fmr1* cohort consisted of 10 KO males and 8 WT litter mate males from the LE-*Fmr1*^*em2Mcwi*^ strain. They underwent two rounds of PPI experimentation, separated by 3 – 4 months, and they were aged 11 – 12 months at the time of the first PPI experimentation and 14 – 15 months at the time of the second PPI experimentation. Between the two experiments, one of the *Fmr1* KO male rats developed a tumor and was euthanized.

The second *Fmr1* cohort consisted of 9 KO males and 9 WT males, all of which were litter mates from the LE*-Fmr1*^*em4Mcwi*^ strain. They underwent one round of PPI experimentation and they were aged 4 – 5 months at the time of the PPI experimentation.

There was one cohort comprised entirely of WT males consisting of 12 rats. They underwent two rounds of PPI experimentation, and were 9 – 11 months at the time of the first experimentation and 13 – 15 months at the time of the second experimentation. As these rats had similar behavioral experiences as the first cohort of *Fmr1* rats, and were experimented on at roughly the same time, we included them in our analysis of the effects of *Fmr1* KO on PPI (Fig. 4). Our conclusions remain unchanged whether or not we included these animals.

The first WT male-female cohort consisted of 6 males and 6 females. They underwent 2 rounds of PPI experimentation separated by 2 months. They were aged 4 – 5 months at the time of the first PPI experimentation and 6 – 7 months at the time of the second PPI experimentation.

The second WT male-female cohort consisted of 6 males and 6 females. They underwent two rounds of PPI experimentation separated by 2 months. They were aged 3 – 4 months at the time of the first PPI experimentation, and 5 – 6 months at the time of the second PPI experimentation.

### Data collection and analysis

Experiments were conducted using four SR-Lab startle systems (San Diego Instruments). The systems were calibrated with a digital sound meter in the center of the test chamber. Each experiment consisted of 12 sessions, with the exception of two experiments that had 6 sessions. On the day prior to the first session, each rat was placed in the apparatus for one hour of constant background sound for initial habituation to the apparatus. Each session began with five minutes of background sound, followed by five habituation trials of a sound 50 dB above the background sound and no prepulse sound. After the habituation trials, sessions consisted of either 5 or 7 repeats of 21 – 48 different stimulus conditions, randomly ordered and separated by inter-trial intervals randomly drawn from the range of 5 – 15 s. Rats completed 2 – 3 sessions per day, and in total, each rat received either 60 or 84 (12 sessions) or 28 – 32 (6 sessions) repeats of each stimulus condition.

A stimulus condition was defined by the three parameters: the startle sound level, the prepulse sound level, and the delay time between the prepulse and startle sounds. The startle sound level varied between 0 – 60 dB above background; the prepulse sound level varied between 0 – 18 dB above background; and the delay time varied between 50 – 200 ms. Prepulse and startle sounds were white noise bursts lasting 20 ms and 40 ms, respectively. The delay time was calculated from the time of prepulse onset. The background sound was either 70 dB or 77 dB, depending on the experiment.

The raw accelerometer readings were first normalized to account for different gains of the startle systems. For each session and rat, we fit a Gaussian distribution to the distribution of accelerometer readings for the first 100 ms of every trial. This is always before the presentation of the startle stimulus, and therefore represents a baseline (Fig. S2a&b). Each accelerometer reading was then z-score normalized by subtracting the mean and dividing by the standard deviation of the Gaussian fit.

Following this normalization, we identified the maximal value within 100 ms following the startle sound for each trial. We then averaged these maximal values across trials at a given condition, which we define as the movement of the animal to that condition. The movement was then used to compute the standard metric for PPI:

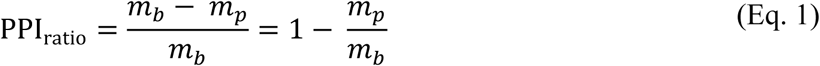

where *m*_*b*_ is the movement of the animal in response to the startle sound alone, i.e. the baseline startle response, and *m*_*p*_ is the movement to the startle sound preceded by a prepulse sound.

We combined the trial repeats of a given condition across all of the sessions of an experiment, since PPI is not thought to habituate across trials^7,24^. Furthermore, we observed only minor changes in baseline startle response between the first and second halves of experiments, such that the mean changes in startle were smaller than the standard deviation between animals, and we did not observe significant between-trial dependencies (data not shown).

### Functional model of PPI

We describe an animal’s baseline startle responses with the equation *m*(s) = *m*_0_ + *N*(s), where *m* is the movement as a function of a startle sound, *s*, and *m*_0_ is the baseline movement independent of sound. We define this equation as the animal’s baseline startle curve, corresponding to startle in the absence of a prepulse sound.

We then introduce scaling parameters to describe how the baseline startle curve is modified by different prepulse conditions. Here, a prepulse condition is defined by the intensity of the prepulse sound and the delay between the prepulse and startle sounds. First, we introduce a parameter, *α*_*c*_, corresponding to the scaling of the startle response due to a prepulse condition, *c*. A model with just startle-scaling is the model that implicitly underlies the traditional PPI_ratio_ metric. After subtracting the baseline movement, *m*_0_, we are left with the following model with just startle-scaling:

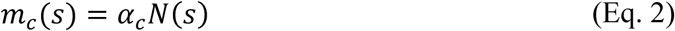

Second, we introduced a parameter, *β*_*c*_, corresponding to the scaling of the startle sound in a specific prepulse condition. This gives us the following model with both startle-scaling and sound-scaling:

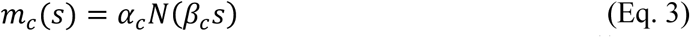

Finally, we used a sigmoid function as the monotonically increasing function, *N*(.), at the basis of our model. This sigmoid describes the specific functional form of the baseline startle curve, i.e. the startle responses without a prepulse sound:

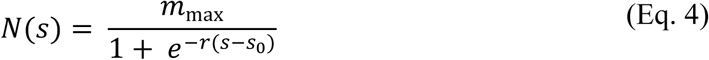

Here, *s* is the startle sound level, *m*_max_ is the maximal movement due to a startling sound, i.e. the saturation point, *s*_0_ is the sound at which the animal startles at 50% of maximal, and *r* is the slope of the sigmoid, describing how rapidly the startle response changes from zero to maximal.

Thus, in total this model contains five parameters: three parameters for the baseline startle curve (*m*_max_, *r, s*_0_) and two scaling parameters for each prepulse condition (*α*_*c*_, *β*_*c*_). The baseline startle curve (Eq. 4) is modified by different prepulse conditions, c, according to the scaling parameters *α*_*c*_ and *β*_*c*_ (Eq. 3). This is a formal model of prepulse inhibition that can be fit to data from individual animals.

We fit all of the data for a given animal with a single fitting routine minimizing the total root mean squared error (RMSE) between the model and the data across all conditions. Initial conditions for the scaling parameters were no scaling (i.e. *α*_*c*_ = *β*_*c*_ = 1 for all *c*); initial conditions for the baseline startle curve were set to the parameters that best fit the baseline data alone, which we obtained by separately fitting a sigmoid to the baseline data. For ease of comparison with the prior PPI metric, the scaling parameters were converted to percentage scaling via 100 * (1 – *α*_*c*_) and 100 * (1 – *β*_*c*_).

We determined whether both startle-scaling and sound-scaling made significant contributions to the fit of the model using a cross-validation approach. Specifically, we cross-validated the model with both startle-scaling and sound-scaling (Eq. 3) and compared against the model with only startle-scaling (Eq. 2) by training each model on 80% of the data and then testing on the remaining 20% of holdout data. For each rat in each experiment, we conducted 100 iterations of cross-validation on randomly selected training and testing data. In each iteration, we computed a normalized RMSE between the models and the testing data, such that the difference between the model and the data at each condition was normalized by the standard error at that condition. Then, for each rat in each experiment, we computed the average normalized cross-validation error across all 100 iterations for both models.

### Group differences in model parameters

The model fits produce five parameters per condition for each animal, and we determined whether groups of animals differed using a linear discriminant analysis (LDA), which finds the hyperplane that best separates the two groups. To visualize this, we projected the data onto the vector orthogonal to the hyperplane, called the linear discriminate (LD), since by definition this is the vector that best separates the groups (Fig. 3b). To evaluate the significance of the LDA, a permutation test on the mean absolute distance from the LDA hyperplane was computed with 10,000 iterations of randomly permuted group labels. In addition, leave-one-out cross-validation was computed for each condition using a permutation test with 10,000 iterations. We define group differences in the model parameters as significant mean absolute distance from the LDA hyperplane (p < 0.05, permutation test) and significant cross-validated classification accuracy (p < 0.05, permutation test) in more conditions than would be expected by chance alone (p < 0.05, bootstrapped ratio test).

### Group differences in PPI

For groups where the LDA analysis revealed a difference, we carried out a second set of analyses to understand the source of the differences. For the comparison between baseline startle parameters and sound and startle scaling, we computed ANCOVAs including a group by baseline interaction term. This interaction term was used to confirm homogeneity of slopes between groups. We also checked for group differences in the baseline parameters using t-tests.

We did not include ANCOVAs for two conditions in which WT male and WT female rats differed in baseline threshold (Fig. 5b), as the ANCOVA is inappropriate in the presence of non-random group differences in the covariate^58^. However, we continued to use ANCOVAs for all other conditions, since, as a whole, the groups did not differ on either baseline covariate. Finally, group effects on startle-scaling and sound-scaling were analyzed using ANCOVAs without a group by baseline interaction term (Fig. 4&5). We define group differences in the sound-scaling or startle-scaling components of PPI as a significant main effect of group (p < 0.05, ANCOVA) in more conditions than would be expected by chance alone (p < 0.05, bootstrapped ratio test).

## Results

We first set out to understand potential causes of inconsistencies in PPI results in the literature. Studies of the *Fmr1* KO mouse report increases^39,49–51,56^, decreases^52,53^, or no difference^54,55^ in PPI compared to WT, and one study concluded that *Fmr1* KO mice show the opposite PPI result compared to humans with Fragile-X Syndrome^39^. As PPI had not been explored in *Fmr1* KO rats, we initially asked whether these inconsistencies could be due to species differences. At the same time, we noted that in the previous studies only a small number (<10) of repeats of any given stimulus condition were used, raising the possibility that variability in PPI measurements also contributed.

We therefore collected data from 28 – 84 (median 60) repeats of each PPI condition in each individual rat. Strikingly, even with the larger number of trials we reproduced the inconsistent results found in mice, both within the same cohort of animals at different sound levels and across different cohorts of animals at similar sound levels. In the first cohort of rats, we varied the prepulse sound and the startle sound, while keeping the delay between the prepulse and the startle sound constant. We found that *Fmr1* KO rats had a lower PPI_ratio_ than WT rats when the startle sound was 30 dB above baseline (p < 0.04, two-way ANOVA) (Fig. 1a). In contrast, *Fmr1* KO rats had a greater PPI_ratio_ than WT rats when the startle sound was 50 dB above baseline (p < 10=G, two-way ANOVA) (Fig. 1a). In the second cohort of rats, we varied the delay between the prepulse and the startle sound, while keeping the prepulse sound constant. We found that *Fmr1* KO rats had a greater PPI_ratio_ than WT rats when the startle sound was 35 dB above baseline (p < 10=H, two-way ANOVA) (Fig. 1b). In contrast, there was no difference in PPI_ratio_ when the startle sound was 50 dB above baseline (p > 0.09, two-way ANOVA) (Fig. 1b), and the trend was in the opposite direction from the 35 dB condition. We found no significant group by prepulse condition interactions (p > 0.05, two-way ANOVA).

**Figure 1.**
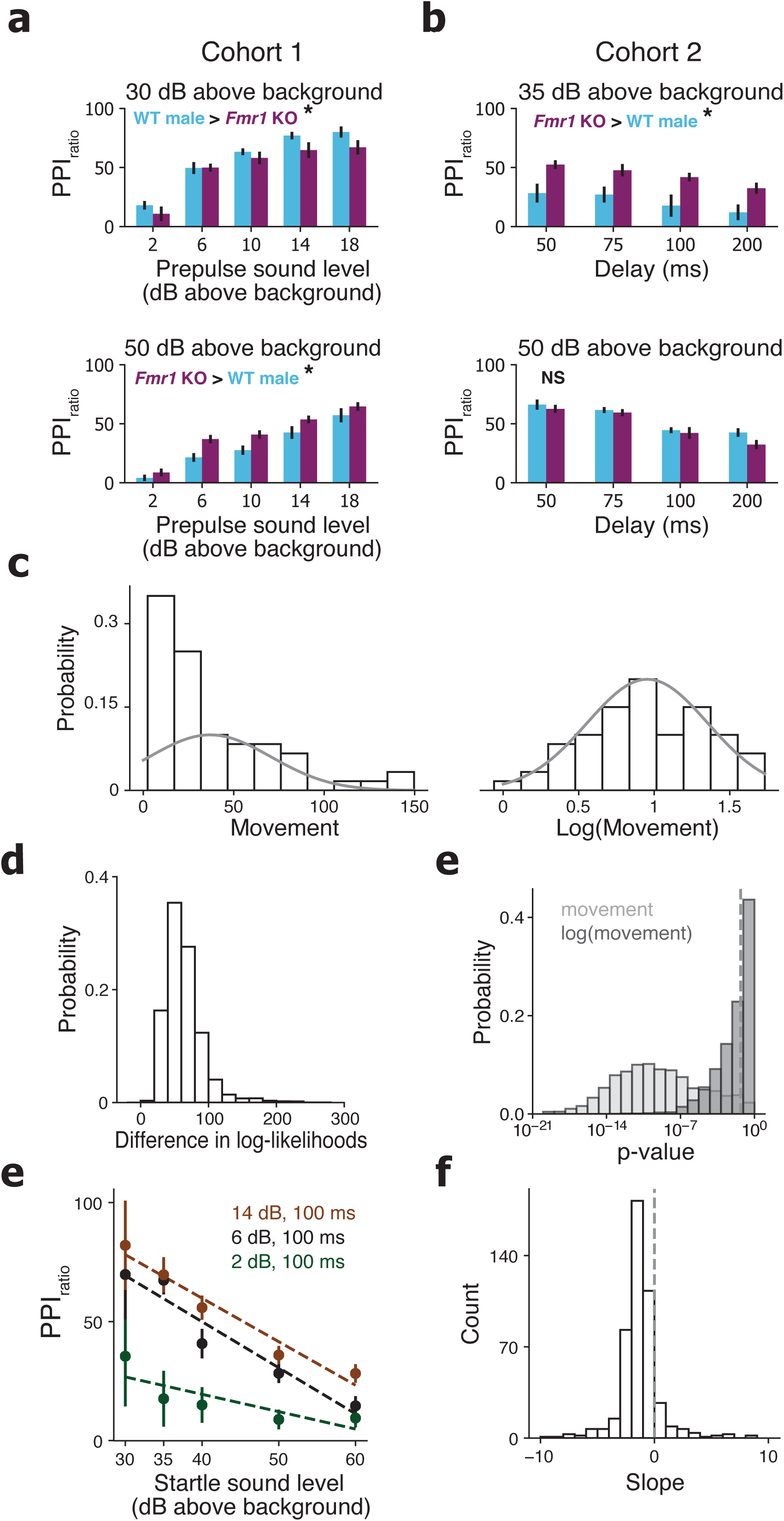
Inconsistencies in standard PPI_ratio_ measurement. (a) PPI_ratio_ from the first *Fmr1* cohort of male rats. Green: WT (n = 7) and red: *Fmr1* KO (n = 9) across different prepulse sounds with a constant 100 ms delay and startle sounds of 30 dB above background (top) or 50 dB above background (bottom). (b) PPI_ratio_ from the second *Fmr1* cohort of male rats. Green: WT (n = 9) and red: *Fmr1* KO (n = 9) across different delays with a constant prepulse of 14 dB above background and startle sounds of 35 dB above background (top) or 50 dB above background (bottom). For a & b, significant group differences were defined as a two-way ANOVA with p < .05. (c) Example probability distribution of gain-normalized movement (see Fig. S2) (left) and log_10_ of gain-normalized movement (right) for one rat to a startle sound of 40 dB above background with no prepulse. Solid curves are Gaussian functions with mean and standard deviation equal to those of the data and height equal to the height of the bin containing the mean. (d) Distribution of the differences in log-likelihood of the movement data under a log-normal distribution and the log-likelihood of the movement data under a Gaussian distribution across all rats and stimulus conditions. (e) Distribution of Shapiro-Wilks normality test p-values across all rats and stimulus conditions for the data before log-transformation (light grey) and after log-transformation (dark grey). Dotted vertical line shows p = .05. Smaller p-values indicate greater probability of rejecting the null hypothesis that the data drawn from a Gaussian distribution. (f) PPI_ratio_ using log-transformed movement data for one rat (same rat as Fig. 1c) across five different startle sound levels (x-axis), three different prepulse sounds (colors), and a constant 100 ms delay. Bars show the standard error of the mean. Dotted lines show linear regressions for each prepulse sound level. (g) Distribution of linear regression slopes of PPI_ratio_ vs startle sound level (dotted lines in Fig. 1f) across all rats and stimulus conditions. Dotted line shows slope of 0.

PPI_ratio_ was also inconsistent between cohorts at similar sound levels. In cohort 1 at 30 dB above baseline, *Fmr1* KO rats had a lower PPI_ratio_ than WT animals (p < 0.04, two-way ANOVA), but in cohort 2 at 35 dB above baseline, *Fmr1* KO rats had a greater PPI_ratio_ than WT animals (p < 10^−4^, two-way ANOVA). In cohort 1 at 50 dB above baseline, *Fmr1* KO rats had a greater PPI_ratio_ than WT animals (p < 10^−3^, two-way ANOVA), but in cohort 2 at 50 dB above baseline, there was no significant difference between *Fmr1* KO and WT animals (p > 0.09, two-way ANOVA), and the trend was in the opposite direction from cohort 1. Thus, we found inconsistent PPI_ratio_ results within and between cohorts, showing that *Fmr1* KO rats exhibit similarly mixed PPI_ratio_ results as seen in *Fmr1* KO mice.

### Invalid assumptions underlie the PPI_ratio_ metric

Previous work identified two additional factors that could contribute to inconsistencies in PPI_ratio_ results: an incorrect assumption of an underlying Gaussian distribution^47^ and an incorrect assumption about the stability of PPI_ratio_ across different startle sounds^48^. Whether these issues are specific to the datasets examined in that past work or more general has not been established. We therefore asked if we could replicate these findings in our cohorts.

Both findings replicated. First, we found that the data are not consistent with an underlying Gaussian distribution but were instead more consistent with a log-normal distribution. (Fig. 1c,d). Across all conditions, only 3.2% of all of the conditions across all animals were consistent with a Gaussian distribution (Fig. 1e) (p > 0.05, Shapiro-Wilks test). In contrast 50.0% of the conditions across all of the animals were consistent with a normal distribution after taking the log of the values, i.e. consistent with a log-normal distribution (Fig. 1e) (p > 0.05, Shapiro-Wilks test). While the log-normal is not a perfect fit, it was a better fit than a Gaussian distribution across all conditions and rats (Fig. 1d), and it represents a good balance between fit and interpretability, so we chose to take the log of the max startle as the basis for our PPI measurements^47^.

Second, we also confirmed that the traditional PPI_ratio_ measure is not the same across different startle sounds, given a constant prepulse condition^48^. If the PPI_ratio_ extracts a core feature of the phenomenon of PPI, then the ratio should be consistent across changes in the denominator, here the startle without a prepulse (Materials & Methods Eq.1). However, we found that not to be the case. Even when using the more accurate log-normal representation of the data, PPI_ratio_ systematically decreases as a function of increasing sound level (Fig. 1f). This decrease was seen in 422 / 488 (86.5%) of prepulse conditions across the 72 rats (Fig. 1g). Thus, the PPI_ratio_ represents only a low-dimensional slice of the phenomenon of PPI, and the ratio does not capture the underlying structure that describes how a prepulse sound modifies the startle response across conditions.

### A new analytical model for PPI

As far as we were able to determine, no formal model underlies PPI_ratio_. However, a very reasonable model for such a ratio would be a scaling of the startle by the prepulse, i.e. *m* = *m*_0_ + *αN*(*s*), where *m* is the movement in response to a startling sound, *m*_0_ is the baseline movement independent of sound, *α* is the startle scaling that occurs due to a prepulse, *s* is the sound level, and *N*(.) is a monotonically increasing function. With such a model, a straightforward derivation (see Supplementary Methods) shows that it is not possible for PPI_ratio_ to decrease with increasing startle sound levels, as long as *m*_0_ ≥ 0 and 0 < *α* ≤ 1. Using this same framework, we developed a model-based analysis of the phenomenon of PPI.

A critical first step was to sample the space of startle responses across many different startle sound levels and many different prepulse conditions, including no prepulse sound. We plotted the startle response as a function of the startle sound and found that, for all of the prepulse conditions, the relationship was well represented by a sigmoid function (Fig. 2a). We therefore chose a sigmoid as the monotonically increasing function at the basis of our model, defined as *N(*.) above (see Materials and Methods). This sigmoid describes an animal’s startle responses to the baseline prepulse condition, i.e. the condition with no prepulse sound. We define this as the animal’s baseline startle curve (Fig. 2a, 0 dB prepulse).

**Figure 2.**
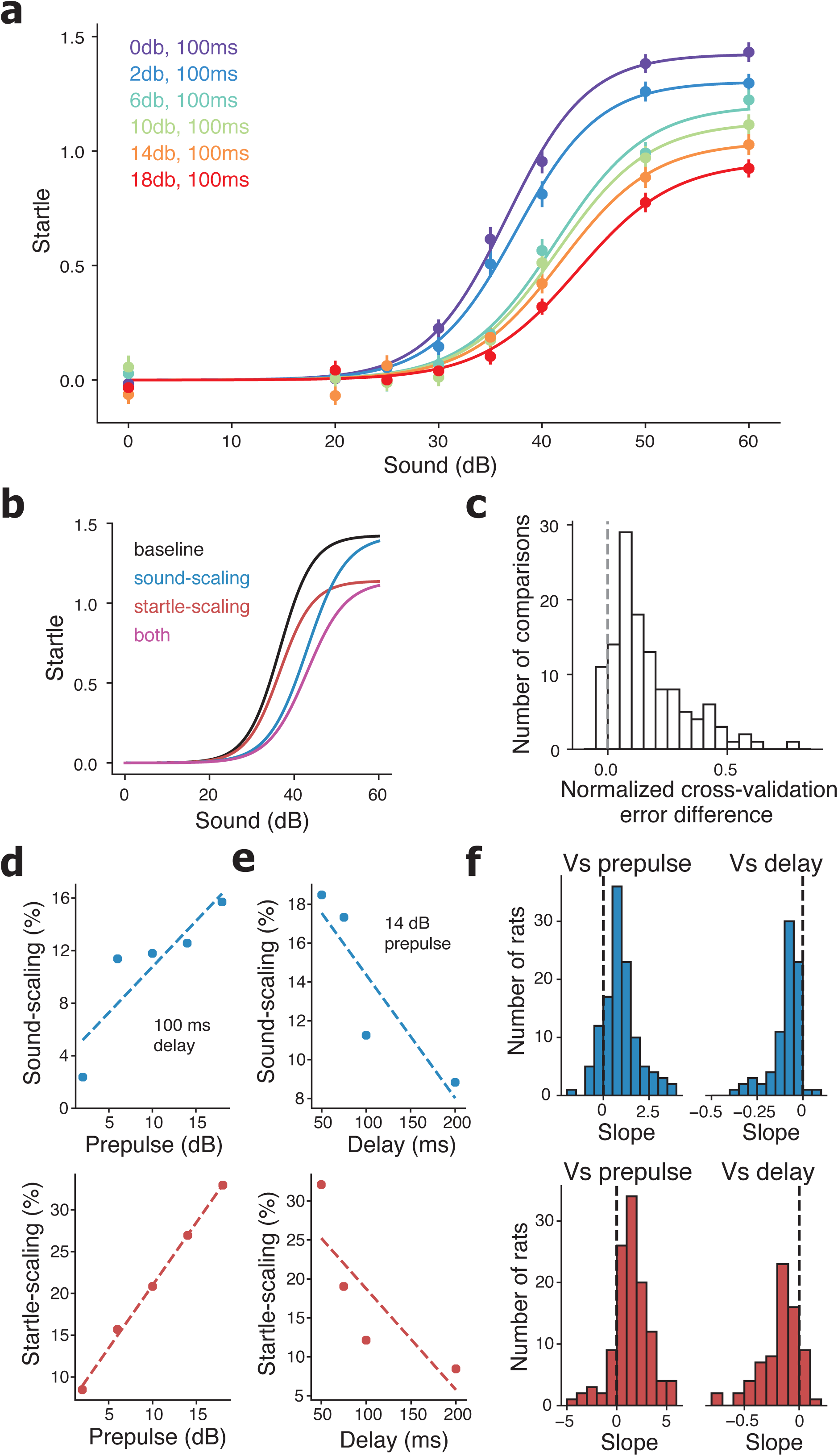
Startle scaling and sound scaling underlie phenomenon of PPI. (a) Startle responses for one rat (same rat as Fig. 1c&f) across different startle sounds (x-axis) and prepulse sounds (colors) with a constant 100 ms delay. Warmer colors indicate louder prepulse sounds. Bars indicate standard error of the mean. (b) Diagram showing how a baseline startle curve (black) could be scaled via sound-scaling (blue), startle-scaling (red), or both startle-scaling and sound-scaling (magenta). (c) Distribution across all rats and experiments of the difference between the normalized cross-validation error of the model with both startle-scaling and sound-scaling and the normalized cross-validation error of the model with only startle-scaling. Positive error differences indicate that the two-parameter model had lower error, and the dotted vertical line shows error difference of 0. (d) Sound-scaling vs. prepulse sound (top) and startle-scaling vs prepulse sound (bottom) for an example rat (same rat as Fig. 2a) from an experiment that varied prepulse level. (e) Sound-scaling vs. delay (top) and startle-scaling vs. delay (bottom) for a different example rat from an experiment that varied delay. For d & e, dotted lines indicate linear regressions. (f) Distribution of linear regression slopes across all rats of sound-scaling vs. prepulse (top left), sound-scaling vs. delay (top right), startle-scaling vs. prepulse (bottom-left), and startle-scaling vs. delay (bottom right). Dotted lines show a slope of 0.

Our next step was to functionally describe how an animal’s baseline startle curve is modified by different prepulse conditions (Fig. 2a). We determined the specific functional form of these modifications to the baseline startle curve by revisiting the interpretation of PPI as one of sensory-motor gating^59^. Sensory-motor gating can occur in two fundamental ways: through modifying the movement that occurs in response to a perceived sound or through modifying the perception of sound. The first modification, startling a different amount in response to the same perceived sound, could manifest through changes in attention^60^ or motor readiness. The second modification, perceiving the same sound differently, could manifest through sensory adaptation^61^.

To disentangle these components, we introduce two parameters, *α*_*c*_ and *β*_*c*_, for each prepulse condition, *c*, which describe how the baseline startle curve is modified by the prepulse condition. Note that a prepulse condition is defined by the prepulse sound level and the delay time between prepulse and startle sounds (see Materials & Methods). For each prepulse condition, *c*, a given perceived sound causes more or less startle as a function of *α*_*c*_, and a given sound is perceived as louder or softer as function of *β*_*c*_. Functionally, *α*_*c*_ and *β*_*c*_ scale the baseline startle curve along the startle and sound axes, respectively, and thereby represent fundamental aspects of the phenomenon of PPI.

This yields the model *m* = *m*_0_ + *α*_*c*_*N(β*_*c*_*s*), where *α*_*c*_ corresponds to startle-scaling and *β*_*c*_ corresponds to sound-scaling at prepulse condition *c*. Note that with *β*_*c*_ < 1 the startle curve expands along the abscissa (sound axis) providing an increase in the difference between the startle curve with a prepulse when compared to without a prepulse. This scaling has the potential to help us understand the observed decrease in PPI_ratio_ with increasing startle sound (Fig. 1f&g): the difference between curves due to differences in sound scaling is maximal near the midpoint and gets smaller as the curves approach their asymptotes (Fig. 2b), which would lead to that decrease.

Our goal was to develop a measure that more accurately reflected the structure of the data as compared to the PPI_ratio_, so we sought to evaluate how well our model fit the data compared to the model that implicitly underlies PPI_ratio_. Namely, our model contains both startle-scaling and sound-scaling, whereas the model that underlies PPI_ratio_ contains only startle-scaling (see Materials & Methods Eq. 1).

We found that our two-parameter model—with both startle-scaling and sound-scaling— had lower cross-validated error than the one-parameter model in 116 / 124 (93.5%) of comparisons (Fig. 2c). Each rat contributed either one or two comparisons, depending on whether the rat was tested in one or two rounds of experimentation. The median normalized error of the two-parameter model was 0.12 lower than the median normalized error of the one-parameter model, meaning that our model with both startle-scaling and sound-scaling was a better fit to the data by ∼12% of the standard error of the data points when compared to the model with just startle-scaling that implicitly underlies PPI_ratio_.

This new model could also explain the known dependencies of PPI_ratio_ on prepulse condition^1,62^ and of self-reported sound intensity on prepulse condition^63,64^. Prepulse conditions with greater magnitude prepulse sounds and shorter delays produced greater scaling of the baseline startle curve (Fig. 2d). To quantify this effect, for each rat we fit lines to the PPI scaling parameters when compared to the prepulse sound intensity (Fig. 2d) and delay (Fig. 2e). We then analyzed the distribution of slopes across all rats, and we found that the distribution mean was significantly nonzero (p < 10^−8^, t-test) (Fig. 2e). This indicates that both sound-scaling and startle-scaling increase with increased prepulse sound intensity and with decreased delay.

Overall, our model greatly reduces the number of parameters required to understand PPI across a range of startle sounds. This is because our model has only two parameters—startle-scaling and sound-scaling—that describe how the animal’s entire baseline startle curve is modified. In contrast, the PPI_ratio_ represents only a slice of the startle curve at a single startle sound level, so many different PPI_ratio_ values would be required to describe the animal’s PPI across a range of startle sounds.

### Analysis of group differences in model parameters

Up until this point we have focused on an accurate understanding of the phenomenon of PPI for each individual animal in each prepulse condition. Specifically, each animal and prepulse condition has five parameters: three describing the baseline startle curve and two describing how PPI scales the baseline startle curve. We next developed a method to determine whether the groups of animals differed in the five-dimensional space of these model parameters.

Given that PPI can be affected by individual-animal factors, such as age^29–31^, experience^43,44^, and strain^27,65^, we restricted our analyses of group differences to comparisons within cohorts of animals whose data were collected at the same time, with animals controlled for age and behavioral experience. We first compared a cohort of animals composed of *Fmr1* KO (n = 18) and litter mate WT males (n = 16) rats, along with a cohort composed of all WT males (n = 12). Separately, we compared the cohorts of animals composed of WT female (n = 12) and WT male (n = 12) rats.

For each prepulse condition, we asked whether the model parameters distinguished between the groups. The five parameters for each animal in each group can be thought of as a point in a five dimensional space, and we therefore used a linear discriminate analysis (LDA) to identify the hyperplane that best separates the points associated with one group from those associated with the other. We then asked whether the mean absolute distance of the points from each group to that plane was greater than expected by chance and whether the cross-validated predictions of group membership were better than chance.

Focusing first on the *Fmr1* KO male and WT male groups, we found no conditions where the animals’ mean absolute distance from the LDA hyperplane was greater than expected by chance (p > 0.05, permutation test)(Fig. 3b, top). We also found that there were no conditions where the cross-validated classification accuracy was greater than expected by chance (p > 0.05, permutation test)(Fig. 3c, left). Thus, the *Fmr1* KO and WT male rats were not separable in their model parameters.

**Figure 3.**
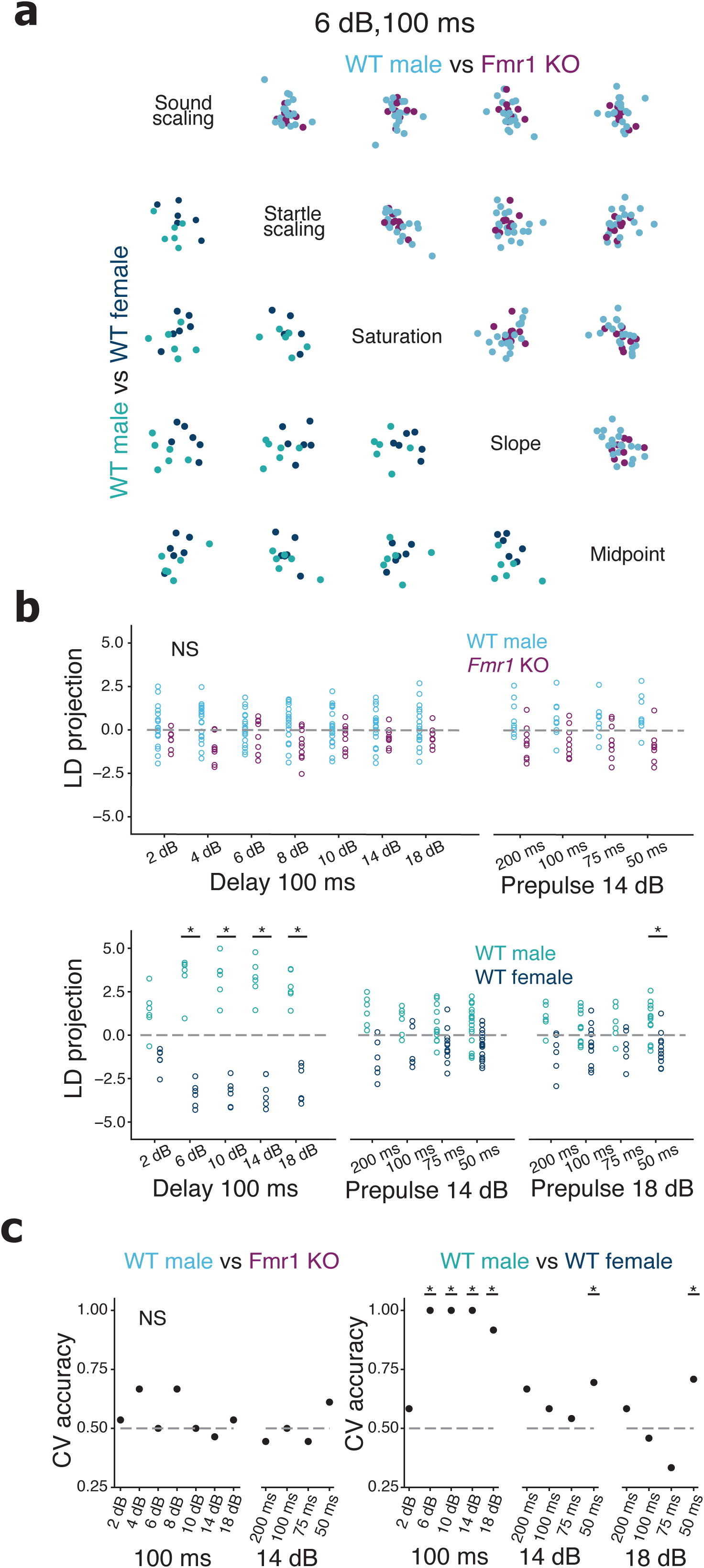
WT male and WT female rats, but not *Fmr1* KO and WT male rats, are separable in their model parameters. (a) Pairwise scatter plots of the model parameters for WT male vs WT female rats (lower left) and *Fmr1* KO male vs WT male rats (upper right) for the condition with a 6 dB prepulse and 100 ms delay. All parameter values are mean-subtracted and standardized between 0 and 1. Main diagonal lists the five model parameters, such that each model parameter defines the x-axis and y-axis of the scatter plots in the same column and row, respectively. (b) Projections of the model parameters for *Fmr1* KO male vs WT male rats (top) and WT male vs WT female rats (bottom) onto the linear discriminate (LD), i.e. the vector orthogonal to the LDA hyperplane that best separates the groups within each prepulse condition. Dashed horizontal lines indicate values that lie on the LDA hyperplane. Asterisks indicate conditions with p < 0.05 for permutation test of total unsigned distance across all rats from the hyperplane separating the groups. (c) Leave-one-out cross validation of LD prediction accuracy for *Fmr1* KO and WT male rats (left) and WT male vs WT female rats (right) for all prepulse conditions. Dashed horizontal lines indicate 50% prediction accuracy. Asterisks indicate conditions with p < 0.05 for permutation test of leave-one-out cross validation accuracy.

Separately, we computed LDA on the model parameters across all rats in the WT male and WT female groups. We found that the animals’ mean absolute distance from the LDA hyperplane was greater than expected by chance in 5/13 prepulse conditions (p < 0.05, permutation test)(Fig. 3b, bottom), which is improbable by chance alone (p < 10^−3^, bootstrapped ratio test). Furthermore, the cross-validated classification accuracy was greater than expected by chance in 6/13 prepulse conditions (p < 0.05, permutation test)(Fig. 3c, right), which is improbable by chance alone (p <10^−4^, bootstrapped ratio test). Thus, the WT female and WT male rats were robustly separable in their model parameters.

### PPI covaries with the baseline startle curve

The results above establish that WT female and WT male rats are different in their responses to the startle sound, but this this does not by itself imply a difference in PPI itself, as the LDA includes both the PPI scaling parameters and the baseline startle parameters. We therefore asked whether our model could also provide insight into potential differences in PPI. To do so we first needed to disentangle the effects of the baseline startle response and of PPI in the observed differences between WT male and female rats. Given that PPI consists of both a scaling of the startle response and a scaling of the perceived sound level (Fig. 2), we asked whether features that describe the baseline startle curve along the startle and sound axes might be covariates of the corresponding PPI scaling parameters.

The baseline saturation is naturally related to startle-scaling as both are in units of startle. The other two baseline parameters—midpoint and slope—describe the relationship between sound level and startle, and jointly they determine an animal’s baseline threshold to startle in the absence of a prepulse. The baseline threshold parameter is naturally related to sound-scaling as both are in units of sound level (Fig. 4a). Thus, we identified four major features of the baseline startle and PPI: two related to the startle axis (startle-scaling and baseline saturation) and two related to the sound axis (sound-scaling and baseline threshold).

**Figure 4.**
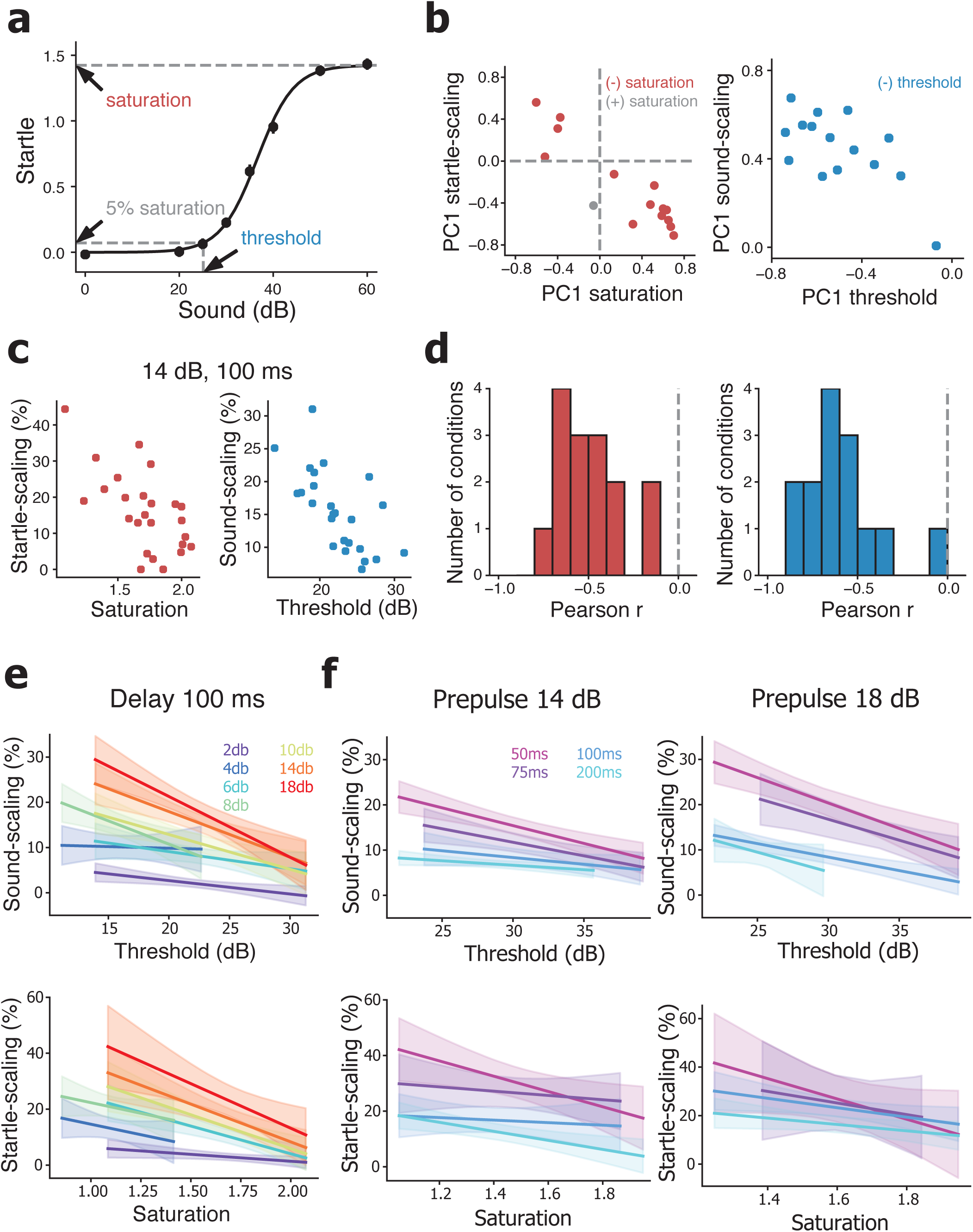
Sound-scaling and startle-scaling covary with baseline startle curve. (a) Example baseline startle curve for the same animal from Figure 2a. Arrows and dashed lines indicate the saturation, defined as the asymptotic maximum of the sigmoid function, and the threshold, defined as the sound at which the curve reaches 5% of saturation. Solid line shows fit of the model to the data. (b) Scatter plot of PC1 startle-scaling weight vs PC1 saturation weight (left) and PC1 sound-scaling weight vs PC1 threshold weight (right) across all prepulse conditions. Dashed vertical and horizontal lines indicate PC1 weights of 0. Red points indicate conditions with opposite direction PC1 weights between startle-scaling and saturation. Blue points indicate conditions with opposite direction PC1 weights between sound-scaling and threshold. (c) Example scatter plots of sound-scaling versus threshold (left, r^2^ = 0.44) and startle-scaling vs. saturation (right, r^2^ = 0.40). For both plots prepulse sound was 14 dB and delay was 100 ms. (d) Distribution of Pearson’s R values for all conditions of sound-scaling vs baseline threshold (left) and startle-scaling vs. baseline saturation (right). Dotted vertical line shows R value of 0. (e) Linear regressions of sound-scaling vs. baseline threshold (top) and startle-scaling vs. baseline saturation (bottom) at different prepulse sound levels. Warmer colors indicate louder prepulse sounds. (f) Linear regressions of sound-scaling vs. baseline threshold (top) and startle-scaling vs. baseline saturation (bottom) at different delays. Warmer colors indicate shorter delays. For e & f, shaded area around lines indicate 95% confidence intervals of the regression.

To understand the structure of these four parameters, we ran principal component analysis (PCA) across all of the WT male animals in each condition. The first principal component (PC1) explained 38% - 70% (mean 52%) of the variance and was significant in 9/15 prepulse conditions (p < 0.05, permutation test)(Fig. S3). This is more conditions than expected by chance (p <10^−6^, bootstrapped ratio test). Strikingly, in 14/15 conditions, the PC1 startle-scaling weight was in the opposite direction of the saturation weight (Fig. 4b, left), and in all 15 conditions the sound-scaling weight was in the opposite direction of the threshold weight (Fig. 4b, right).

These opposing signs suggest a relationship between sound-scaling and threshold, and separately, between startle-scaling and saturation. Indeed, within each prepulse condition, we found that startle-scaling was negatively correlated with the saturation level of the baseline startle curve across all of the WT male rats (Fig. 4c&d). Animals with higher startle saturation, i.e. higher maximum startle, tend to have less startle-scaling. The mean Person’s r was −0.49 ± 0.05, and the r^2^ values ranged from 0.02 to 0.62 (Fig. 4d), meaning that the startle saturation accounted for up to 62% of the variance of the startle scaling across rats within prepulse conditions. This correlation was significant in 8/15 conditions.

Similarly, within each prepulse condition, we found that sound-scaling was negatively correlated with startle threshold of the baseline startle curve across all of the WT male rats (Fig. 4c&d). Animals with higher startle thresholds tend to have less sound scaling. The mean Pearson’s r was −0.60 ± 0.05, and the r^2^ values ranged from 0 to 0.83 (Fig. 4d), meaning that the startle threshold accounted for up to 83% of the variance of the sound scaling across rats within prepulse conditions. This correlation was significant in 10/15 conditions.

These results imply that we can only interpret PPI with respect to individual animals’ baseline startle parameters. We therefore computed linear regressions for startle-scaling as a function of baseline saturation and for sound-scaling as a function of baseline threshold (Fig. 4e&f). The PPI scaling vs. baseline correlations showed a range of values (Fig. 4d) where the slope of the regressions increased with increasing prepulse sound level and with shortened delay (Fig. 4e&f and S4). Thus, the phenomenon of PPI is both a function of prepulse condition as well as of the baseline startle curve.

### Analysis of PPI group differences

These findings indicated that we could determine whether differences between the groups were due to differences in PPI by adjusting for baseline covariates. For each prepulse condition, we computed two linear regression models across all of the WT male rats and, separately, across all of the WT female rats. These linear models describe the two PPI vs. baseline correlations: sound-scaling vs. baseline threshold (Fig. 5a&b) and startle-scaling vs. baseline saturation (Fig. 5c&d).

**Figure 5.**
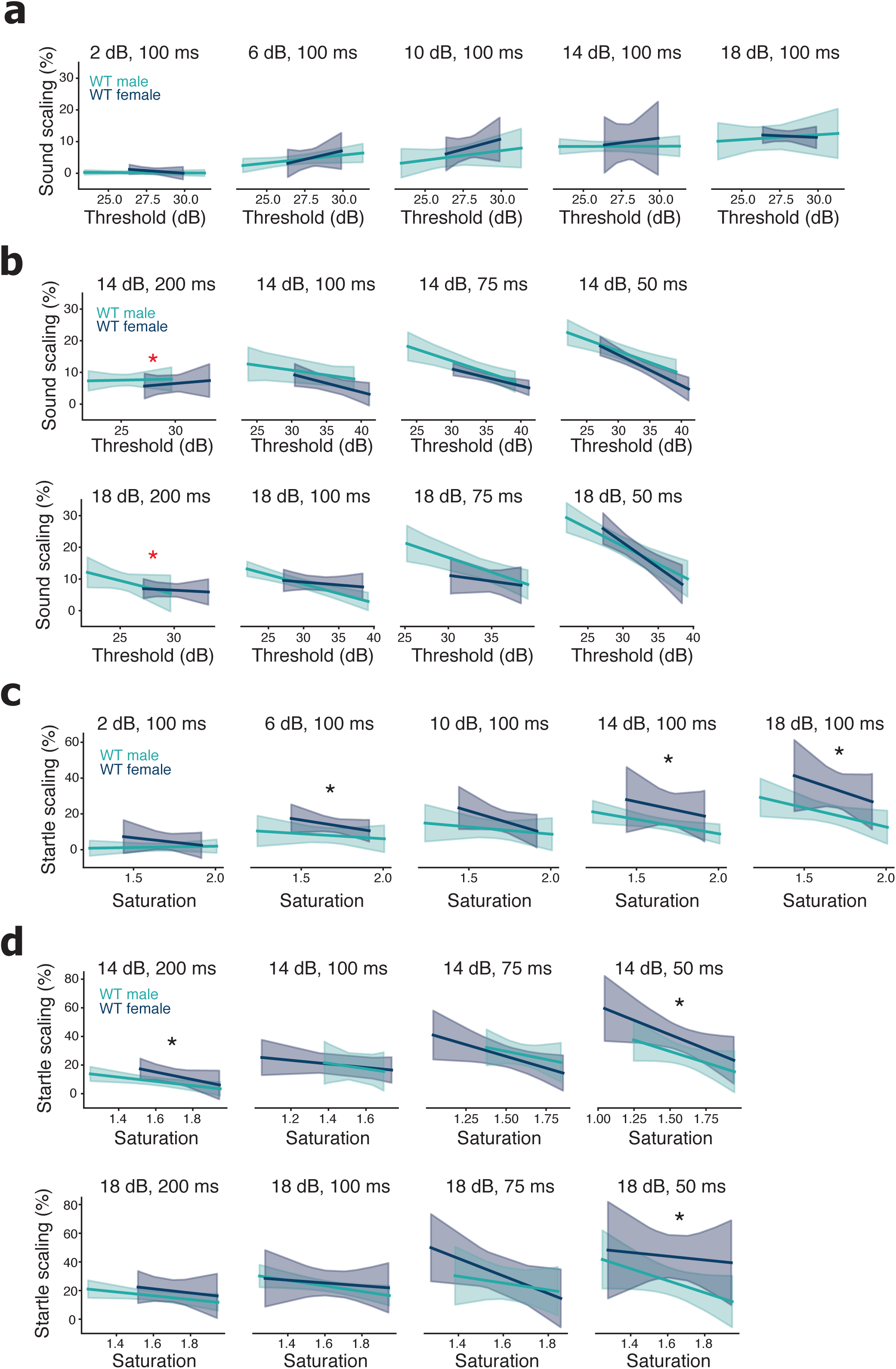
Less startle-scaling in WT male than WT female rats. (a) Linear regressions of sound-scaling vs. baseline threshold for WT male (green) and WT female (grey) rats from the single experiment that varied prepulse sound level. Subplots show increasing prepulse sound level from left to right. (b) Linear regressions of sound-scaling vs. baseline threshold for WT male and WT female rats from the three experiments that varied delay. Subplots show decreasing delay from left to right. Red stars indicate conditions with a group difference in baseline threshold (p < .05, t-test). This was not significant after controlling for multiple comparisons (p > .05, bootstrap ratio test), but these two conditions were excluded from ANCOVAs. (c) Linear regressions of startle-scaling vs. baseline saturation for WT male and WT female rats from the single experiment that varied the prepulse sound level. Subplots show increasing prepulse sound level from left to right. (d) Linear regressions of startle-scaling vs. baseline saturation for WT male and WT female rats from the three experiments that varied delay. Subplots show decreasing delay from left to right. For c & d black stars indicate conditions where WT male rats had lower startle-scaling than WT female rats (p < .05, ANCOVA). For a, b, c, & d there were no baseline by group interactions (p > .05, ANCOVA). In total, WT male rats had lower startle-scaling than WT female rats in 6/13 conditions, which is significant after multiple comparisons (p < .05, bootstrap ratio test).

Next, for each prepulse condition, we computed an ANCOVA with interaction term. After confirming that there was no baseline by group interactions (p > 0.05), we re-ran the ANCOVAs without an interaction term. We found no conditions with a group difference in baseline saturation (p > 0.05, t-test), but we did find two conditions with a group difference in baseline threshold (Fig. 5b) (p < 0.05, t-test). However, a control for multiple comparisons reveals 2/13 significant conditions to be insignificant (p > 0.1, bootstrapped ratio test), and the exclusion of those two conditions did not affect the results. Thus, the differences between groups could not be explained by a difference in baseline parameters.

We then considered the effects of group on PPI. We found that WT female rats had greater startle-scaling than WT male rats (p < 0.05, ANCOVA) in 6/13 conditions, which is significant after controlling for multiple comparisons (p <10^−4^, bootstrapped ratio test). In contrast, we found no differences between WT female and WT male rats in sound-scaling at any condition (p > 0.05, ANCOVA). As a confirmation, running LDA on the parameters of these models—startle-scaling, sound-scaling, saturation, and threshold—results in 7/13 significant conditions, including the same 6 significant conditions as found with ANCOVA (data not shown).

Finally, to confirm our previous results, we carried out the same analysis on the *Fmr1* KO and WT male rats. There were no conditions where the *Fmr1* KO male rats differed from WT male rats in either startle-scaling or sound-scaling (p > 0.05, ANCOVA)(Fig. S5), nor in either baseline saturation or baseline threshold (p > 0.05, t-test)(Fig. S5). This confirms our finding that *Fmr1* KO male rats were not separable from the WT male rats in their model parameters (Fig. 3b&c). These results in the *Fmr1* KO rat provide a reliable approach that could be used to clarify the inconsistent PPI results with *Fmr1* KO mice and humans with Fragile-X Syndrome.

## Discussion

We found inconsistent PPI_ratio_ results in *Fmr1* KO rats across different startle sound levels within and between cohorts (Fig. 1a&b), reproducing the inconsistent results seen in the *Fmr1* KO mouse literature^39,49–56^. Furthermore, we confirmed that the acoustic startle response is better described by a log-normal than a Gaussian distribution (Fig. 1c,d,&e) and that the PPI_ratio_ changes across startle sound levels^48,66^ (Fig. 1f&g). These results reveal important limitations of the standard PPI methodology.

To address these limitations, we developed a new model of PPI (Fig. 2a), which describes how a prepulse sound scales the baseline startle curve along both the startle and sound axes (Fig. 2b). We found that our model was a consistently better description of the data for individual animals than the implicit model underlying the PPI_ratio_ metric (Fig. 2c). We then found that *Fmr1* KO male rats were not separable from wild-type controls in their model parameters. In contrast, we found that WT male and WT female rats are linearly separable in their model parameters (Fig. 3).

Seeking to explain these differences, we found that startle-scaling and sound-scaling were correlated with the baseline startle response curve across animals within conditions (Fig. 4b – d). Taking this into account, we analyzed group differences in startle-scaling and sound-scaling by fitting linear models to the scaling versus baseline data. We found no difference in PPI between *Fmr1* KO and WT rats (Fig. S5). We did, however, find that WT female rats showed greater PPI startle-scaling than WT male rats (Fig. 5c&d). These findings were robust to changes in startle sound level, and they were reliable across different cohorts of animals.

### Benefits of a new model of PPI

The phenomenon of PPI exists independent of the PPI_ratio_ metric or any other model used to describe it. Fundamentally, animals tend to startle less when a startling stimulus is preceded by a weak prepulse, compared to when a startling stimulus is presented alone. But what is the functional form of this phenomenon, and how does it depend on the stimulus condition and individual differences between animals? The usefulness of PPI in neuroscience and psychiatry depends on our ability to understand the phenomenon itself, and this in turn depends on the metrics used to describe the phenomenon.

Here we present a novel model that disentangles two components underlying PPI: sound-scaling and startle-scaling. In contrast, the model that implicitly underlies the PPI_ratio_ metric does not describe scaling of the startle sound. Previous work has observed changes in perceived sound after a prepulse^63,64,67^, but this has been conceptualized as a separate phenomenon from PPI, often measured using self-report scales. Our model unifies startle-scaling and sound-scaling, revealing them to be two components of the PPI phenomenon, both of which can be observed in the acoustic startle data.

Furthermore, both sound-scaling and startle-scaling are biologically interpretable. Sound-scaling could manifest through rapid sensory adaptation in auditory hair cells^68^ or higher auditory pathways^69^, while startle-scaling could manifest through changes in attention and concentration or other cognitive or motor factors^60^. A formal model that separately parameterizes sound-scaling and startle-scaling allows for a principled deconstruction of the behavioral neurobiology of PPI. In our case, we were able to quantify how much of each parameter contributed to PPI in individual animals (Fig. 2), how they covaried with the baseline startle (Fig. 4), and how they compared between groups (Fig. 5, S5).

If PPI is to be a useful biomarker of disease^20^ or a predictor of treatment outcomes^21^, then at a minimum we need to describe the core behavioral features of the phenomenon using a reproducible methodology. By showing that startle-scaling and sound-scaling underlie the phenomenon of PPI, our model provides such a methodology.

### Assumptions and limitations of the model

One of the challenges that we faced in deconstructing the current way in which the phenomenon of PPI is measured was in disentangling the many assumptions that underlie the current metric used to describe the phenomenon. Therefore, we feel it crucial to lay out the assumptions, and the potential limitations of those assumptions, that underlie our proposed model. We used many repeats of each stimulus condition, which is important given the high variability of the startle response between trials. The alternative—using a small number of trial repeats—suffers from a potentially inaccurate representation of the underlying startle distribution.

However, combing data across many trials assumes that the startle response is relatively stable across trials. Furthermore, by randomly presenting dozens of different stimulus conditions within a session, we assume that there are minimal between-trial dependencies. We did not find evidence to overturn these assumptions, and PPI is not thought to habituate across trials^7,24^. Therefore, we chose to combine data across trials within an experiment, allowing us to develop an accurate statistical representation of the acoustic startle response and a reliable static model of PPI. Further experiments and inquiry could incorporate a more dynamic picture into the interpretation of the phenomenon of PPI, using the model proposed here as a foundation.

As we have justified (Fig. 1c – e), the basis for our measurement of the startle response rests upon the assumption of the log-normality of the data. Skewed data in complex biological systems is a common finding^70^, reflecting interactions in complex systems such as the brain. However, further experimentation could expose that the startle distribution could be more accurately described by more complex distributions, such as variants of the gamma distribution or combinations of several distributions. For example, it is possible that a scalar measurement of startle magnitude only makes sense in a subset of “true startle” events, as distinct from “no startle” events. If so, it could be useful to consider a probability of startle in addition to the magnitude of startle.

We also introduced a new axis along which PPI changes an animal’s response to a startle sound: by scaling the perception of sound itself. To start with the simplest possible model, we assumed that the sound scaling occurred through a single parameter that multiplied the sound axis. It is also possible that there is an additional parameter that shifts the sound axis, and further experimentation will be necessary to determine its role in the phenomenon of PPI.

There is more to PPI than just a ratio of the startle with a prepulse to the startle without a prepulse. PPI is a complex phenomenon which depends on many features of the stimulus and which shows high variability between individual animals. In spite of this complexity, the extensive literature linking PPI_ratio_ to schizophrenia^4–9^ and other disorders^10–19^ suggests that PPI could be a useful methodology for generating mechanistic insights into neuropsychiatric disease. As a step toward that goal, our analytical model allows for a deconstruction of the underlying structure of PPI, which in turn enables robust and replicable studies of the neural circuits underlying PPI and how those circuits vary among individuals in the context of disease.

## Acknowledgements

We thank N. Swerdlow and T.O. Sharpee for helpful discussions and P.W.E. Spratt and A.K. Gillespie for technical assistance. This work was supported by the Simons Foundation Autism Research Initiative (L.M.F. and M.R.D.); by a Jane Coffin Childs Memorial Fund for Medical Research postdoctoral fellowship, and an NIH R25 (R25MH060482) (D.B.K.).

## Author contributions

D.B.K. designed the study; D.B.K. and E.A.M. performed the experiments; D.B.K., E.A.M., and L.M.F. developed the analyses; D.B.K. and E.A.M. analyzed the data; A.M.G., M.N.G., and M.R.D. developed and initially characterized the *Fmr1* knockout rats; and D.B.K., E.A.M., A.M.G., and L.M.F. wrote the manuscript.

---

**Supplemental Figure 1.**
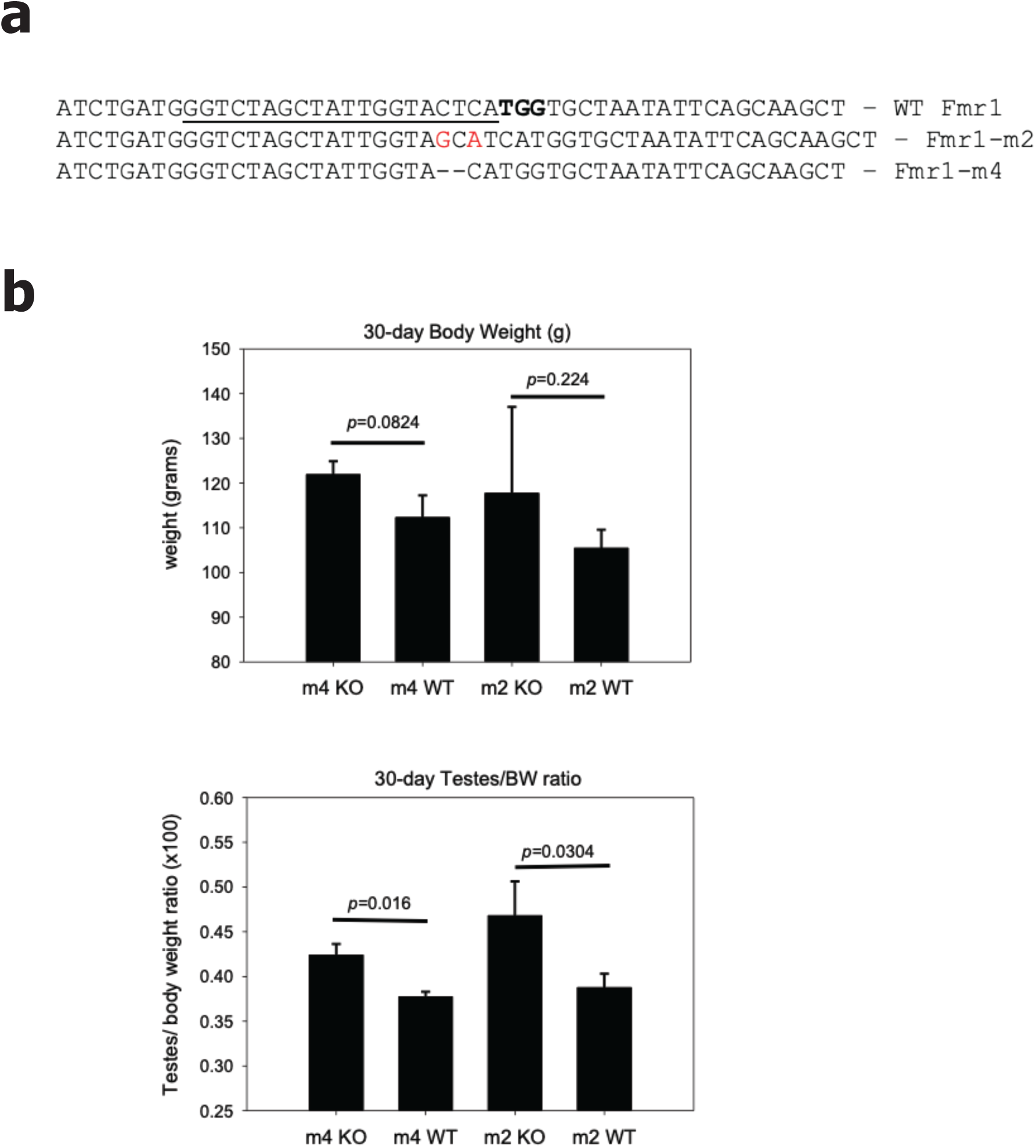
Characterization of Fmr1 knockout rats. (a) Two mutant models were generated (Fmr1-m2 and Fmr1-m4) with frame-shift indels in exon 7. CRISPR-SpCas9 target site is underlined with protospacer adjacent motif (PAM) in bold. (b). Body weight trended larger, and testes/body weight ratio was greater in m2 and m4 knockout males compared to wildtype littermates at 30 days of age. N=3-5 per group, significance determined by Student’s T-test.

**Supplemental Figure 2.**
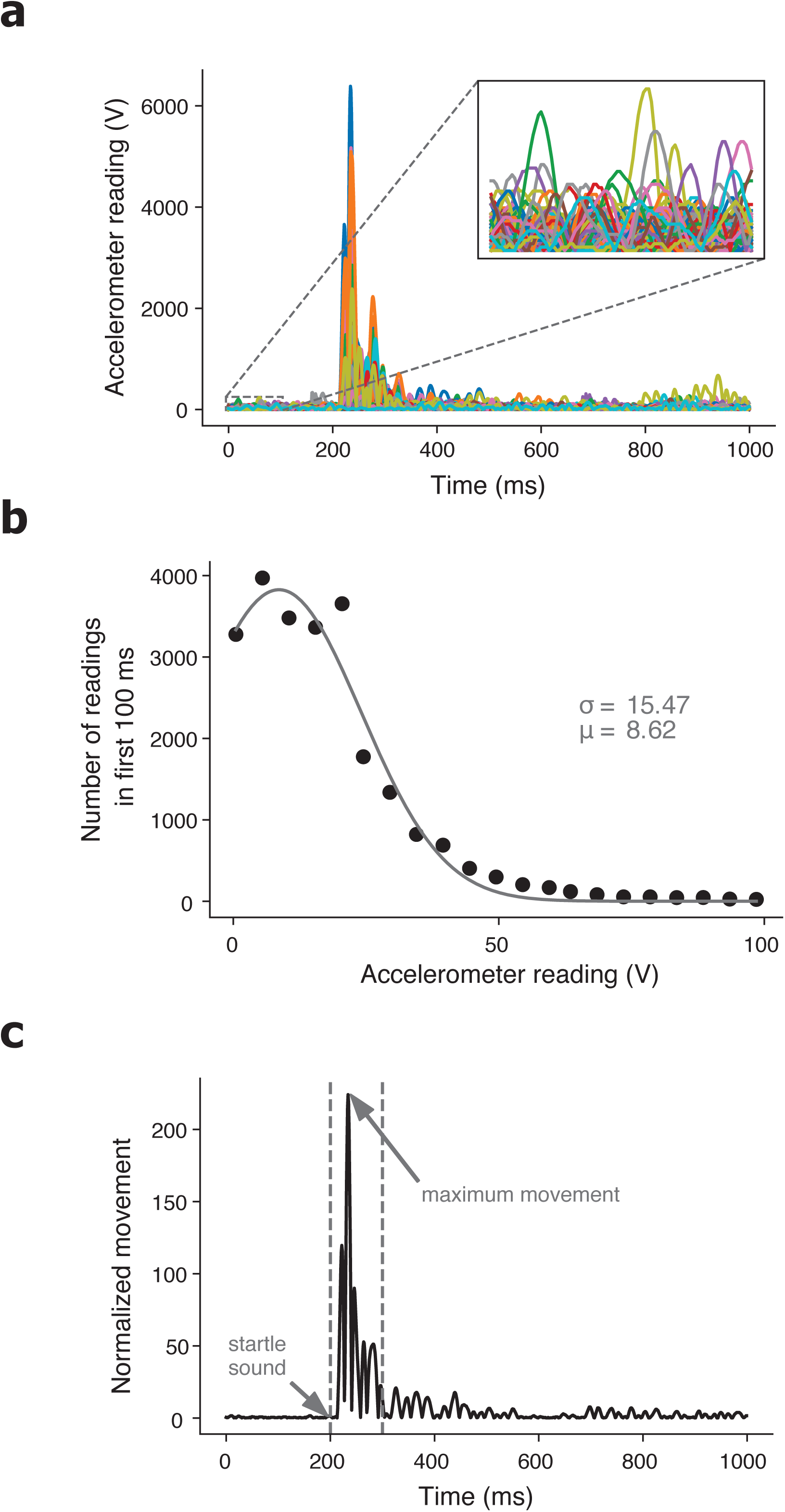
Normalizing movement data for apparatus gain. (a) Raw accelerometer data for all trials (colors) in a single session for one rat. Inset shows accelerometer data in the first 100 ms of the trials. (b) Histogram of accelerometer readings in the first 100 ms across all trials in a single session for one rat (same rat as Fig. S2a). Solid curve shows the Gaussian fit to the histogram, and the legend shows the mean and standard deviation of this Gaussian. (c) Normalized movement of a single trial for one rat (same rat is Fig. S2a&b). Dashed vertical lines indicate the startle sound onset (left) and 100 ms after the startle sound onset (right). Arrows indicate startle sound onset and the maximum normalized movement in the 100 ms window.

**Supplemental Figure 3.**
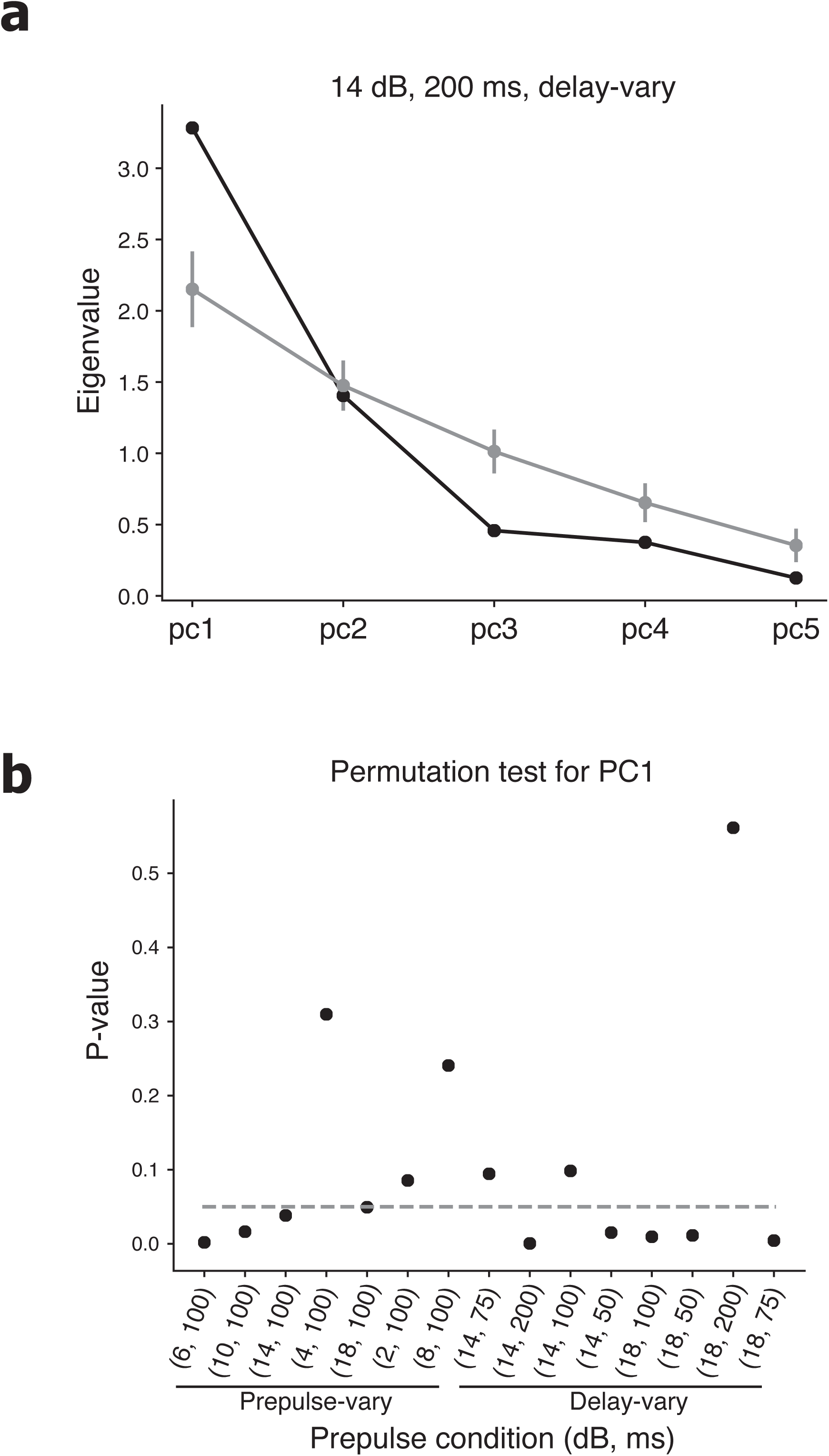
First principal component captures significant variance in the model parameters. (a) Eigenvalues of the five principal components (PCs) of the reduced set of model parameters (startle-scaling, sound-scaling, saturation, and threshold) across all WT male rats for the condition with 14 dB prepulse and 200 ms delay. Grey curve shows mean and standard deviation of the eigenvalues across random permutations of the parameters. (b) P-values of the first PC eigenvalue from a permutation test on the parameter values, across all prepulse conditions. Dashed horizontal line indicates p = 0.05.

**Supplemental Figure 4.**
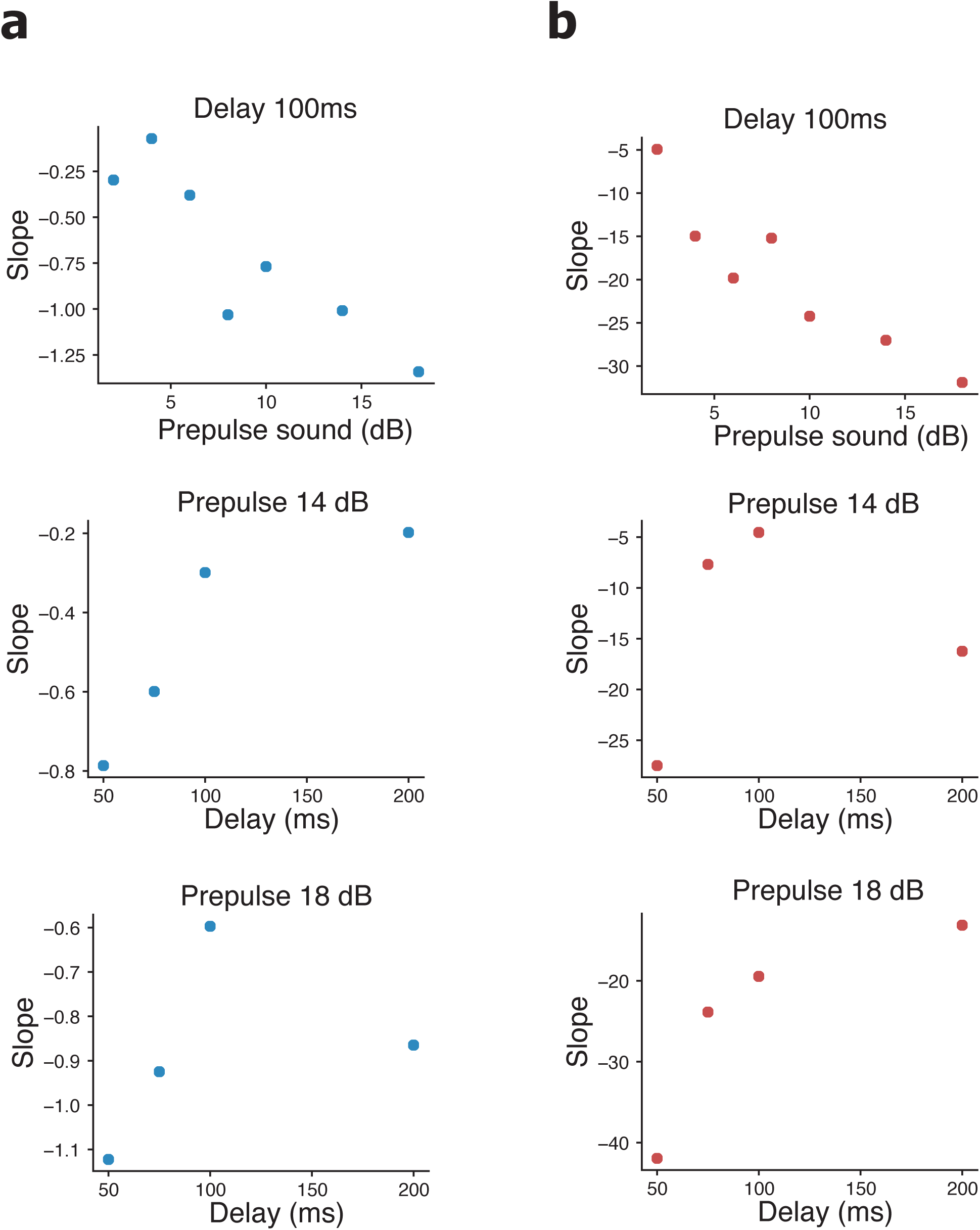
PPI is more sensitive to differences in baseline startle with louder prepulse sounds and shorter delays. (a) Slope of the sound-scaling vs baseline threshold regression slopes as a function of prepulse sound level at a constant 100 ms delay (top); as a function of delay at a constant prepulse level of 14 dB above baseline (middle); and as a function of delay at a constant prepulse level of 18 dB above baseline (bottom). (b) Slope of the startle-scaling vs baseline saturation regression slopes as a function of prepulse level at a constant 100 ms delay (top); as a function of delay at a constant prepulse level of 14 dB above baseline (middle); and as a function of delay at a constant prepulse level of 18 dB above baseline (bottom).

**Supplemental Figure 5.**
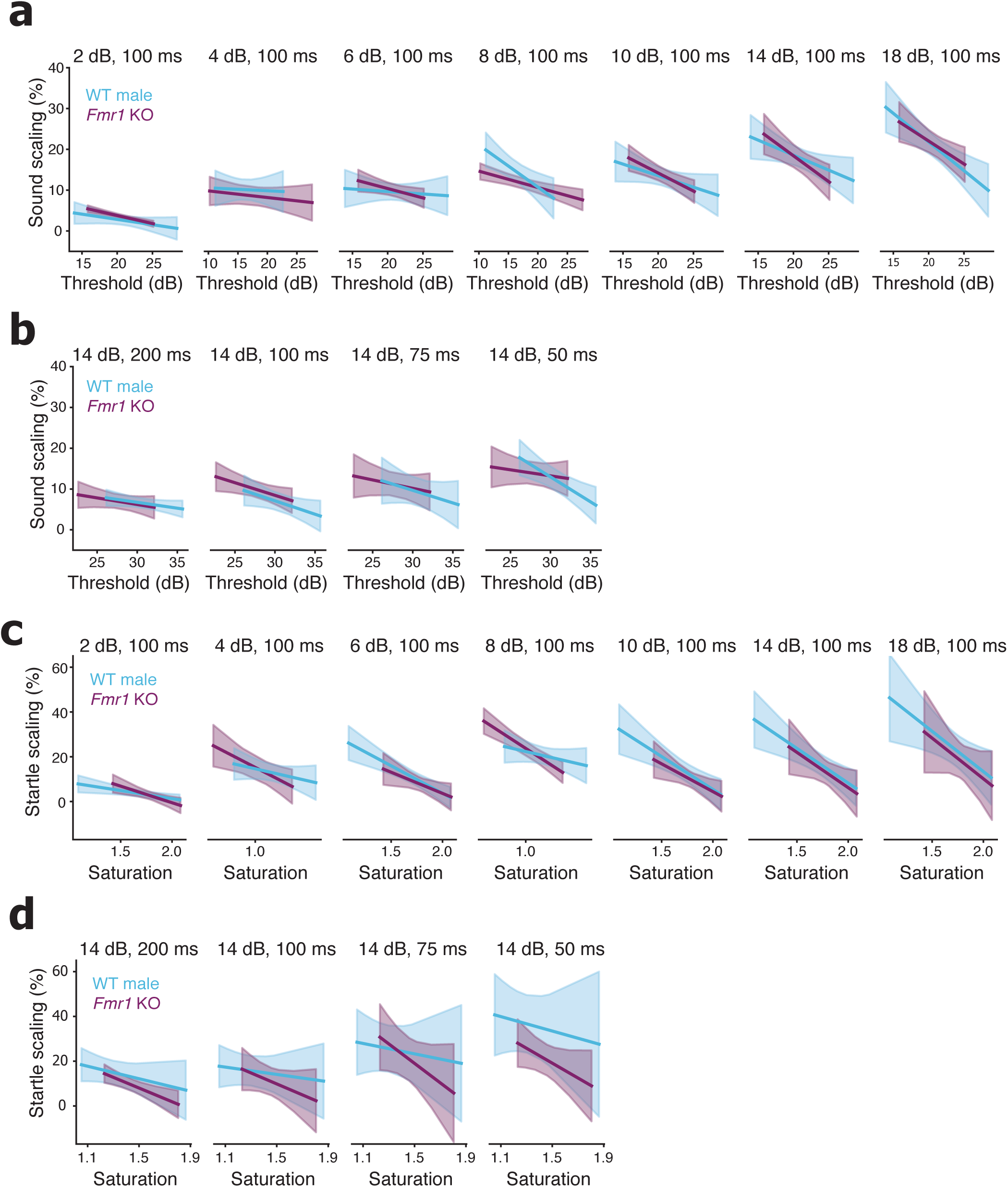
No difference in startle-scaling or sound-scaling between *Fmr1* KO and WT male rats. (a) Linear regressions of sound-scaling vs. baseline threshold for *Fmr1* KO male (red) and WT male (green) rats from experiments that varied the prepulse sound level. Subplots show increasing prepulse sound level from left to right. (b) Linear regressions of sound-scaling vs. baseline threshold for *Fmr1* KO male rats and WT male rats from experiments that varied delay. Subplots show decreasing delay from left to right. (c) Linear regressions of startle-scaling vs. baseline saturation for *Fmr1* KO male rats and WT male rats from experiments that varied the prepulse sound level. Subplots show increasing prepulse sound level from left to right. (d) Linear regressions of startle-scaling vs. baseline saturation for *Fmr1* KO male rats and WT male rats from experiments that varied delay. Subplots show decreasing delay from left to right. For a, b, c, & d there were no group differences in baseline parameters (p > .05, t-test), no baseline by group interactions (p > .05, ANCOVA), and no group differences startle-scaling or sound-scaling at any condition (p > .05, ANCOVA).

## Supplemental methods

Here we derive that the PPI_ratio_ metric can never decrease as a function of increasing startle sound if PPI is just due to a scaling of the startle response, under the assumption that the acoustic startle response is well captured by any monotonically increasing function.

Assume for the sake of contradiction that PPI_ratio_(*s*_1_) > PPI_ratio_ (*s*_1_) for some startle sounds *s*_1_ < *s*_2_. By the definition of PPI_ratio_ (Materials & Methods Eq. 1), this gives:

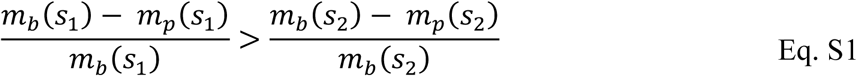

where *m*_*b*_ is the average startle response to the startle sound alone, i.e. the baseline startle response, and *m*_p_ is the average startle response of the animal to the startle response preceded by the prepulse sound.

Let *N* (*s*) be any monotonically increasing, strictly positive function of the sound that well-describes the acoustic startle response. Furthermore, let *m*_0_ be a constant non-negative y-offset due to baseline movement. Therefore, *m*_*b*_ (*s*_1_) = *m*_0_ + *N*(*s*_1_) is the baseline startle response at sound *s*_1_, and *m*_*p*_(*s*_1_) = *m*_0_ + *αN*(*s*_1_) is the startle response at sound *s*_1_ following a prepulse, where *α* is the startle-scaling parameter. Plugging this into Eq. S1, this gives us:

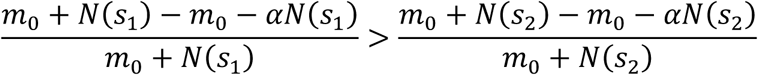

The left side of the equation calculates the PPI_ratio_ metric at sound *s*_1_ and the right side of the equation calculates PPI_ratio_ metric at sound *s*_2_. We then subtract off the right side of the equation and cancel out the baseline movements (*m*_0_) in the numerators such that:

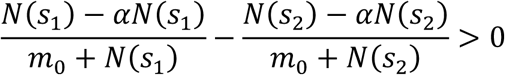

If *α* ≠ 1, then we can then factor out 1 – *α* from both numerators and divide them out of the equation. Furthermore, by definition, PPI implies that 1 – *α* > 0, so we maintain the direction of the inequality, giving:

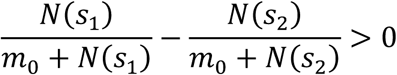

We then combine the fractions and simplify the numerator to get:

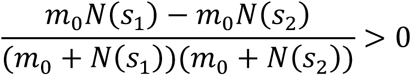

Assuming that the baseline movement, *m*_0_, is strictly positive, so we can factor *m*_0_ out from the numerator:

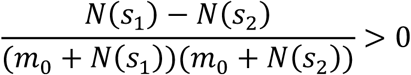

This cannot be true given our definition of 4(0) as a monotonically increasing function that is always greater than zero. Thus, we have reached a contradiction, and Eq. S1 must be false.

Thus, the PPI_ratio_ metric can never decrease if PPI is just due to scaling the startle response, and if PPI is due only to a scaling of the startle response, under the assumption that the acoustic startle response is well captured by any monotonically increasing function that is always positive.

### Protocol for measuring and fitting sound and startle scaling PPI model

1. Measure the acoustic startle response at many different conditions.
  a. Vary the startle sound level across the full range of values over which the startle response changes. For example, we varied the startle sound level between 0 - 60 dB above background, as this was the range over which our rats’ startle responses varied from zero to maximum startle.
  b. Vary the prepulse sound level and/or the delay time across the range of values over which PPI changes as a function of that parameter. For example, we varied the prepulse sound between 0 – 18 dB above background and the delay between 50 – 200 ms.
  c. For each stimulus condition—i.e. combination of prepulse sound, delay time, and startle sound—collect data from at least 50-100 trial repeats for every animal tested.
  d. For each trial, normalize the raw accelerometer data by a baseline accelerometer measure, e.g. by the data from times prior to the presentation of any stimulus (Fig. S2a&b).
  e. Take the log_10_ of all of the normalized accelerometer data.
  f. For each trial, find the maximum value of the log normalized data in a 100 ms window following the presentation of the startle sound.
2. Compute the average startle at each stimulus condition
  a. For each rat, find the mean and standard error of the trial maxima from 1e at each stimulus condition.
  b. For every mean startle value from 2a, subtract the mean value from all of the control conditions, i.e. the conditions with startle sound level 0 dB above background across all prepulse conditions.
  c. We define the resulting values as the startle to a given condition for an animal, and we can plot these values as startle response vs. startle sound (Fig. 2a).
3. Fit the PPI model to the average startle data for each rat
  a. Implement a sigmoid function with startle-scaling and sound-scaling parameters (Materials and Methods Eq. 3&4). This function should accept 5 parameters: *m*_max_, *s*_0_, and A for the baseline sigmoid and *α* and *β* for the scaling due to a prepulse.
  b. Implement an objective function that computes the *total* RMSE error between the average startle response data and the model under a set of parameter choices for *every* prepulse condition. This objective function should take the following parameters: the set of average startle data, the baseline sigmoid parameters (*m*_max_, *s*_0_, and *r*) and the set of all scaling parameters (*α*_c_ and *β*_c_ for all prepulse conditions *c*).
  c. Use a minimization algorithm (e.g. Scipy.optimize) to find the optimal model parameters that minimize the objective function against the average startle data for each rat. Initial conditions for the scaling parameters can be set to no scaling. Initial conditions for the baseline sigmoid can be set to anything that you think will optimize the chances of converging on the best fit.
4. Evaluate group differences in the model parameters
  a. Standardize the parameters to all range between 0 – 1 and subtract the means.
  b. For each prepulse condition, run a linear classifier such as linear discriminate analysis (LDA).
  c. Compute the mean absolute (unsigned) distance from the linear discriminate hyperplane.
  d. Compute LDA classification accuracy using leave-one-out cross-validation.
  e. Report group separability if the mean absolute distance and the cross-validated classification accuracy are greater than expected by chance from permutation tests on the group labels.
5. Find baseline threshold and saturation for each animal
  a. The baseline saturation is defined as *m*_max_ of the baseline sigmoid for a given animal.
  b. Compute the baseline threshold, defined as the startle sound level at which an animal’s baseline startle curve reaches 5% of *m*_max_.
6. Evaluate group differences in PPI
  a. For each prepulse condition, fit two linear models per group: one for sound-scaling vs. baseline threshold and one for startle-scaling vs baseline saturation, and plot these with 95% confidence intervals (Fig. 4&5).
  b. For each prepulse condition, check for group difference in the baseline parameter. If there are significant group differences in the baseline parameter, an ANCOVA cannot be computed for that condition. You can run t-tests for group difference in the scaling parameters but be aware that these differences could be caused by non-random group differences in the baseline startle.
  c. Assuming no/few group differences in the baseline parameters, compute two ANCOVAs for each prepulse condition: one for startle scaling as a function of group and baseline saturation, and one for sound scaling as a function of group and baseline threshold. Include a group by baseline interaction terms in all ANCOVAs.
  d. If the baseline by group interaction terms are significant in any of the ANCOVAs, we cannot use those conditions because they break the homogeneity of slopes assumption.
  e. Assuming no/few significant interaction terms, recompute all of the ANCOVAs without interaction terms, and look for significant main effects of group.
  f. Control for multiple comparisons, where each of your prepulse conditions is a separate comparison, using a bootstrapped ratio test to determine the probability of seeing a given number of significant conditions by chance alone. Alternatively, control for multiple comparisons using Bonferroni correction or related methodology.
  g. Report group differences in PPI startle-scaling or sound-scaling if it holds up to the control for multiple comparisons.

